# Multi-Mode Fiber-Based Speckle Contrast Optical Spectroscopy: Analysis of Speckle Statistics

**DOI:** 10.1101/2022.10.25.513089

**Authors:** Chen-Hao P. Lin, Inema Orukari, Christopher Tracy, Lisa Kobayashi Frisk, Manish Verma, Sumana Chetia, Turgut Durduran, Jason W Trobaugh, Joseph P. Culver

## Abstract

Speckle contrast optical spectroscopy/tomography (SCOS/T) provides a real-time, non-invasive, and cost-efficient optical imaging approach to mapping of cerebral blood flow. By measuring many speckles (n>>10), SCOS/T has increased signal-to-noise ratio relative to diffuse correlation spectroscopy, which measures one or a few speckles. However, the current free-space SCOS/T designs are not ideal for large field-of-view imaging in humans because the curved head contour cannot be readily imaged with a single flat sensor and hair obstructs optical access. Herein we evaluate the feasibility of using cost-efficient multi-mode fiber (MMF) bundles for use in SCOS/T systems. One challenge with speckle contrast measurements is the potential for confounding noise sources (e.g., shot noise, readout noise) to contribute to the standard deviation measure and corrupt the speckle contrast measure that is central to the SCOS/T systems. However, for true speckle measurements, the histogram of pixel intensities from light interference follows a non-Gaussian distribution, specifically a gamma distribution with non-zero skew, whereas most noise sources have pixel intensity distributions that are Gaussian. By evaluating speckle data from static and dynamic targets imaged through MMF, we use histograms and statistical analysis of pixel histograms to evaluate whether the statistical properties of the speckles are retained. We show that flow-based speckle can be distinguished from static speckle and from sources of system noise through measures of skew in the pixel intensity histograms. Finally, we illustrate in humans that MMF bundles relay blood flow information. © 2022 Optica Publishing Group

---

Cerebral blood flow (CBF) is an important biomarker of brain health; however, current imaging techniques are restricted by issues such as lack of bedside capability, ionizing radiation, high cost, and/or low signal-to-noise ratio (SNR) [1]. The brain consumes ~20% of the human body’s energy even though it makes up only ~2% of the body’s mass [2]. Therefore, the brain must receive a substantial amount of the body’s blood supply to maintain this high energy consumption. Consequently, the brain has developed several mechanisms to control CBF to meet its needs, including autoregulation and neurovascular coupling [3,4]. The failure of these mechanisms and the subsequent dysregulation of CBF are hallmarks of many diseases, including traumatic brain injury, stroke, and brain tumors. The gold standard for non-invasively mapping CBF in the clinic is positron emission tomography (PET) [5]. PET has whole-brain coverage and good spatial resolution; however, PET has high instrumentation costs, uses ionizing radiation, and cannot image continuously at the bedside [1]. Arterial spin labeling magnetic resonance imaging (ASL-MRI) is an alternative to PET increasingly used in the clinic [6]. ASL-MRI has whole-brain coverage, great spatial resolution, and does not use ionizing radiation, but it also cannot be used continuously at the bedside and has high instrumentation costs [1]. A low-cost alternative to PET and MRI is diffuse correlation spectroscopy (DCS) [1]. DCS is an optical modality that can be used for continuous bedside monitoring without ionizing radiation; however, DCS has low SNR and only offers a spot measurement of CBF [7]. Thus, clinicians would benefit from a method of measuring CBF that addresses these concerns. Speckle contrast optical spectroscopy/tomography (SCOS/T) methods provide a potential solution to these challenges [7,8]. In this paper we evaluated whether cost-efficient MMF bundles can be used to relay speckle dynamics reflecting the fluid dynamics of a sample, whether statistical properties of the speckles are retained, and whether flow-dependent speckle contrast measures can be distinguished from noise-based, with the overall goal of confirming that MMF bundles are suitable for fiber-based SCOS/T.

Free-space SCOS/T has been introduced as a low-cost optical modality for imaging CBF at the bedside with an improved signal-to-noise ratio (SNR) relative to a typical DCS implementation [9]. SCOS/T techniques estimate fluid flow by measuring the speckle contrast in images from many fluctuating speckles formed by constructive and destructive interference. While DCS can have larger SNR for a single speckle, SCOT increases SNR by combining hundreds to thousands of speckles simultaneously [10].

In free-space SCOS/T, laser light is raster-scanned across the surface of the brain and the diffuse light is measured at a distance by a camera detector. The free-space SCOS/T systems have been shown to obtain similar results as fMRI for relatively large ischemic stroke cores in rodents [11]. However, free-space SCOS/T is not ideal for imaging a large area of the human brain because the contour of the head is not readily captured by a single imaging sensor, and hair occludes the field of view. In contrast, optical fibers can comb through hair to reach the scalp and can be arranged to follow complex head geometries [12]. And a single imaging sensor can be used to collect light from a fiber array with a complex arrangement on the head as long as the sensor ends of the fibers are arranged in a single focal plane for relaying light to the imaging sensor [13,14] With intensity measures only, fiber-based high-density diffuse optical tomography (HD-DOT) systems have been established for imaging the visual cortex and through hair [14].

Single-mode or few-mode fibers are often considered for building fiber-based speckle contrast imaging systems. Single-mode fibers (SMFs) are preferred as light detectors in DCS systems to capture the temporal dynamics of single speckles. Previously, our group has demonstrated that SCOS with few-mode fibers is feasible for relaying flow information [15]; however, few-mode coherent fiber bundles (Schott, IB1050760CTD) suffer from high fragility and prohibitive cost (~$6000 per bundle). To address these limitations, here we explore the feasibility of measuring speckle contrast with multi-mode fiber (MMF) bundles, made of individual fibers with areas 100x-1000x the size of individual speckles. Intuitively, this makes some sense, as MMF has been used for more than a decade for illumination with DCS systems. However, the feasibility of using MMF as detectors in a SCOS/T system has not been well studied. Recent research indicates that pulsatile flow can be measured through individual MMFs [16,17], but the measurement statistics have not been reported to confirm flow-based speckle statistics. There are conceptual reasons to be skeptical about SCOS/T detection via MMF, as the phase field of the light from the surface will be scrambled when the light field transmits through a multi-mode fiber that has an area 100x to 1000x the size of a speckle.

To evaluate the feasibility of using MMF for SCOS, we started with a design for a one-source, one-detector SCOS system, which would serve as one of many channels in a hypothetical SCOT system. The potential advantages of the larger area of the MMFs, compared to SMFs, are higher integrated photon flux and the transmission of many more speckles for greater SNR [18,19]. The potential worry is that somehow the speckle statistics needed for accurate speckle contrast measurements could be corrupted due to mode mixing or other sources of variance in the pixel measurements. In this study, we aimed to determine whether speckle, speckle statistics, and speckle contrast measures from dynamic media can be distinguished from potential confounding noise sources (e.g., detector shot noise).

Our fiber-based SCOS system (Fig. 1a) uses a laser diode (Thorlabs, λ = 785 nm, LP785-SAV50) as a light source and an sCMOS camera (Andor Technology Zyla 5.5, pixel size 6.5µm) as the detector. One end of the MMF bundle (NA = 0.66, 50 µm core-diameter fibers), placed in contact with the sample, captured and relayed the light patterns from the sample to the detector. The detector end of the fiber bundle was imaged on the sensor plane using an objective lens (Mitutoyo, NA = 0.42). The magnification of the imaging system is 20X, resulting in a speckle-to-pixel size ratio of 6.8, generously satisfying Nyquist sampling requirements.

**Fig. 1.**
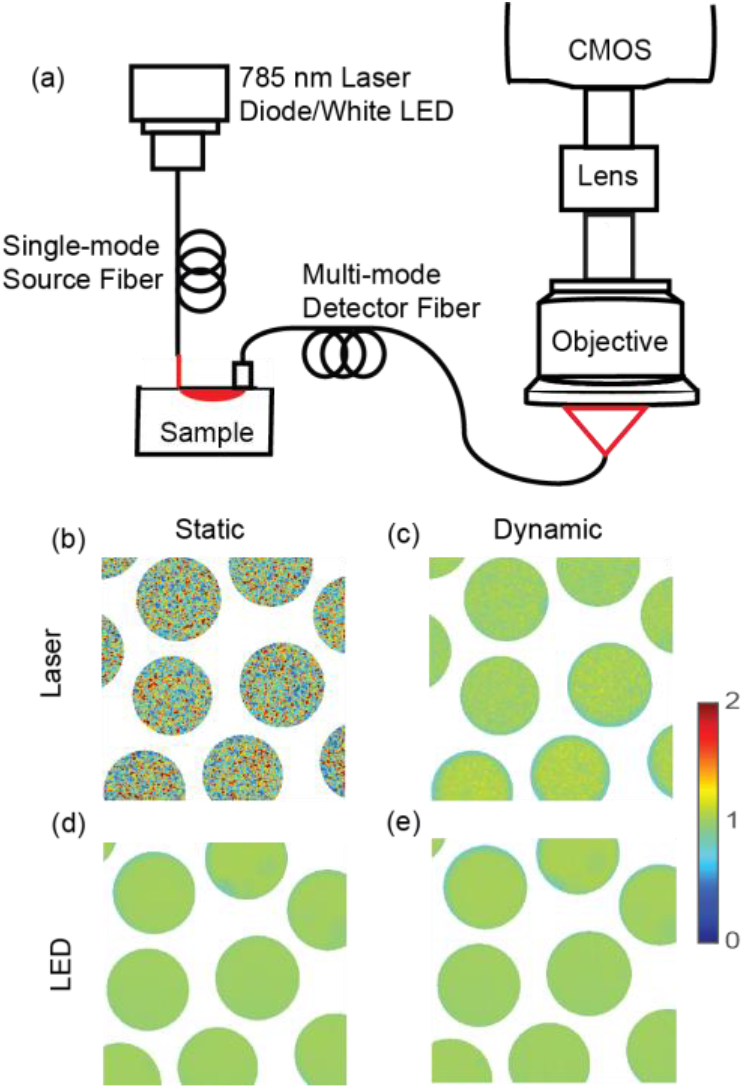
Intensity images of static (silicone) and dynamic (milk) phantoms illuminated by laser or LED with a source-detector distance of 4 mm and detected via multi-mode fiber (MMF) bundles (50 μm core diameter) under long (100 ms) exposure. The system schematic is shown in (a). A granular speckle pattern is apparent when coherent laser light illuminates a static silicone phantom (b), but not a dynamic milk phantom (c) where the decorrelation time of speckle is shorter than the exposure time. Non-coherent LED light illumination also shows no speckle pattern for either the static (d) or dynamic (e) phantoms. These experiments demonstrate that MMF bundles can be used to detect speckle patterns from a sample.

To demonstrate that MMFs can be used for detecting speckle patterns and that the granular patterns arise from coherent interference patterns, we began by manipulating both the illumination and the phantoms (Fig. 1). To manipulate the illumination, we compared an incoherent (LED) source with that of a coherent source (laser diode). The LED source was broadband white light, and the laser was a stabilized single-frequency laser diode. We also varied the sample to compare a static phantom (silicone gel-like tissue phantom), in which the speckle pattern should not change, with a dynamic phantom (whole milk), in which the speckle pattern should move.

We compared intensity images of static and dynamic phantoms illuminated by laser or LED and detected via an MMF bundle under a long (100 ms) exposure time. Under laser illumination, we observed the expected granular speckle patterns for the static phantom. In contrast, for the dynamic phantom, we measured smooth and low fluctuation intensity patterns. These results agree with the basic expectations of speckle theory that motion of the scatterers within the liquid phantom caused the speckle pattens to move within the exposure time (100 ms) resulting in a temporal blurring and lower variance. As a negative control, patterns of both static and dynamic phantoms illuminated by LED were smooth and did not vary between phantoms. These experiments provide a simple but direct demonstration that MMF bundles relay speckle patterns in a way that is sensitive to fluid dynamics.

To further validate the use of MMF we manipulated the exposure times. Speckle contrast images (Fig.2 a-d) were generated directly from the raw intensity images. For each pixel, speckle contrast was computed over a spatial window as the ratio of the standard deviation and mean of the intensity, K = σ/<I>. High speckle contrast is observed for the static phantom with both short and long exposure time but only with the short exposure time for the dynamic phantom. In contrast, low speckle contrast is observed for the dynamic phantom with a long exposure time (Fig. 2d). These results are consistent with dynamic phantoms producing low speckle contrast when the exposure time is long enough for the speckle patterns to move significantly.

**Fig. 2.**
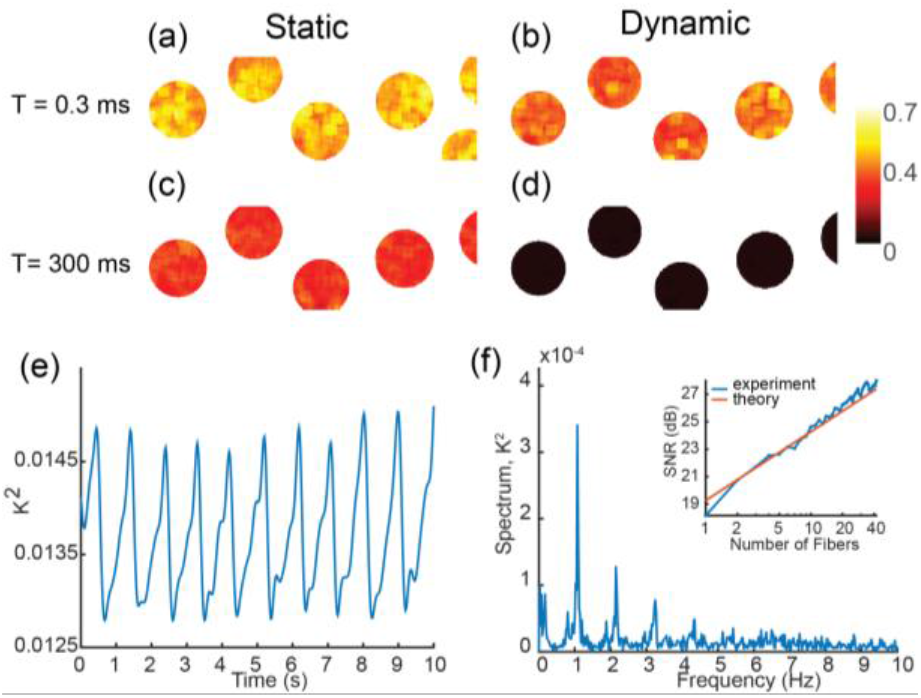
Experiments demonstrating that MMF bundles can be used to relay flow information. Speckle contrast depends on the decorrelation time of the sample and the exposure time of the camera, with long decorrelation times for static phantoms and short decorrelation times for dynamic phantoms. Speckle contrast images derived from the intensity images demonstrate that speckle contrast measurements reflect flow in dynamic media that is not present in static media, with short (a) and long (c) exposure times in static phantom and short (b) and long (d) exposure times in the dynamic phantom. With long exposure time, low speckle contrast seen for the dynamic phantom is due to blurring caused by motion. In vivo speckle contrast waveforms (e) computed from measurements (source-detector distance of 26 mm) on the scalp of a human subject show expected pulsatile flow. The magnitude spectrum (f) shows several harmonics of the pulse frequency, and SNR (f, inset) varies as expected 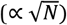 as speckle contrast is averaged over multiple fibers from the MMF bundle.

To confirm that speckle measurements are sensitive to blood flow in tissue directly, we used our system to measure speckles from the scalp of a human subject (Fig. 2e). Speckle contrast computed from measurements (26 mm source-detector distance, magnification of 10X) showed pulsatile flow, with spectral signal-to-noise ratio (SNR) of the pulse peak increasing, up to 27 dB (inset Fig. 2f) as K is averaged over additional fibers from the MMF bundle.

Since speckle contrast is a measure of random fluctuations, we also considered the impact of other sources of variance, in particular experimental noise sources such as dark noise or readout noise. To compare the impacts of measurement noise and flow, we collected intensity data from the static and dynamic phantoms with and without illumination. We generated histograms (Fig. 3) for data collected either temporally or spatially and computed statistics (Table 1) of these intensity distributions (Fig. 3 a,b). We fit the data with Gamma and Gaussian distributions and used a Kolmogorov-Smirnov (KS) test to determine goodness of fit, i.e., rejecting the suitability of a distribution if the p value is < 0.05 (which indicates the fit statistically differs from the data). Conceptually, while data from a Gaussian distribution will be fit well by a gamma function with skewness ~0, data with a non-Gaussian distribution will only fit well to a gamma distribution, with a measurable positive skew. Dark noise measurements produced symmetric histograms in both temporal and spatial analysis and are well fit by a Gaussian distribution - as expected. For the static phantom, temporal analysis produced a symmetric distribution that was also well fit by a Gaussian function with low skewness. In contrast, the spatial analysis of the static phantom produced a right-skewed gamma distribution, with a poor Gaussian fit indicative of true speckle statistics. In contrast to the static phantom, both spatial and temporal analysis of the dynamic phantom data produced right-skewed histograms, because the speckles are moving in time. Overall, the skewness statistic and the related shape parameter of the gamma distribution distinguished between signals coming from noise versus optical scattering speckle patterns. To further test whether MMF bundles alter speckle distributions, we compared speckle images of the static phantom measured directly (no MMF) with images measured through an MMF bundle. We found similar statistics, with skewness of 1.48 directly vs 1.42 with the bundle.

**Fig. 3.**
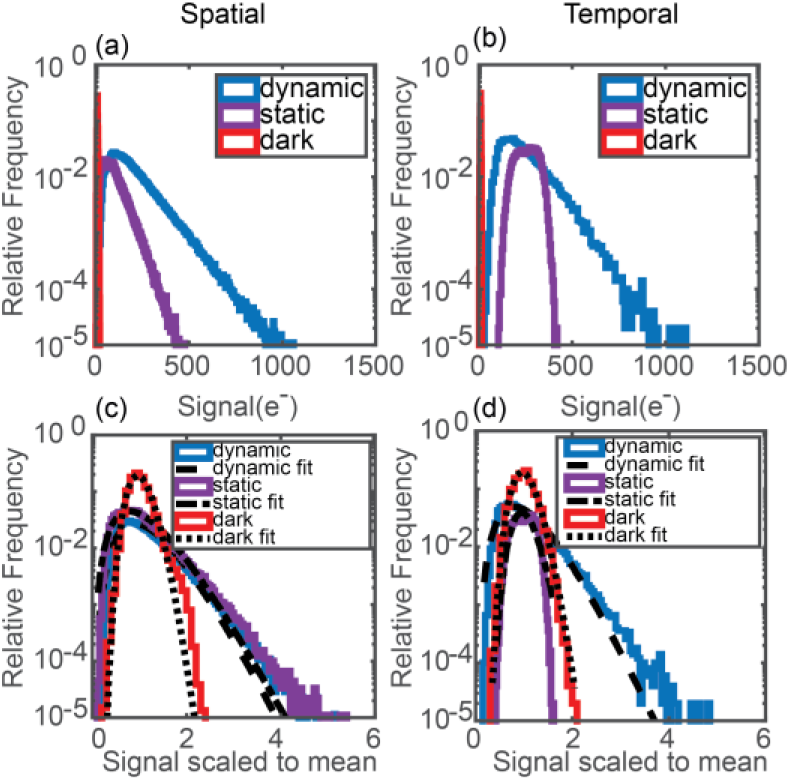
Statistical distributions differentiate speckle signals from measurement noise. Spatial histograms were created from single frames of speckle images, and temporal histograms were created from a time series at a very high frame rate (5411 fps) for a single pixel. Spatial (a) and temporal (b) histograms are shown for dynamic and static phantoms imaged with laser illumination, and for a non-illuminated, control condition (dark noise only, static phantom). Histograms of intensities normalized to the mean within each fiber were fit with gamma distributions (spatial, c, temporal, d). The skewness was found to distinguish noise (low skewness) from speckle (high skewness).

**Table 1.** Statistics of speckle signal and noise.

|  |  | Real speckle signals |  |  | Noise |  |  |
| --- | --- | --- | --- | --- | --- | --- | --- |
|  |  | Dynamic spatial | Dynamic temporal | Static spatial | Static temporal | Dark spatial | Dark temporal |
| Intensity Data | Mean, $\mu$ (e-) | 212 | 229 | 105 | 258 | 9.32 | 12.2 |
| | Std, $\sigma$ (e-) | 114 | 108 | 54.8 | 53.2 | 2.0 | 2.13 |
|  | Skew | 1.7 | 1.35 | 1.45 | -0.0898 | 0.945 | 0.0441 |
| Gamma | Shape parameter | 4.21 | 5.07 | 4.15 | 22.3 | 22.9 | 31 |
|  | KStest p value | 0.109 | 0.593 | 0.563 | 0.912 | 0.892 | 0.999 |
| Gaussian | KStest p value | 0.00175 | 0.0428 | 0.0138 | 0.912 | 0.674 | 0.999 |

The spatial sampling rate of speckle images can affect the quality of speckle contrast data. To determine the impact of sampling on flow-related intensity statistics such as skewness, we synthetically controlled the sampling rate by downsampling our images (Fig 4). We investigated the effect of sampling on the integrity of the probability density function (PDF) of the intensity and showed that the skewness can be used to determine whether the speckles are sampled properly. Speckle sizes were obtained using the Gaussian fit of auto-covariance and scaled to the airy disk. Histograms from speckle signals are right-skewed when they are measured properly, which is usually considered when it satisfies the Nyquist sampling theorem (speckle size should be at least two times the pixel size). We demonstrated that the intensity distribution has the desired skewness only for speckle/pixel size ratios > 1.7, while for speckle/pixel < 1.7 there is inadequate sampling.

**Fig. 4.**
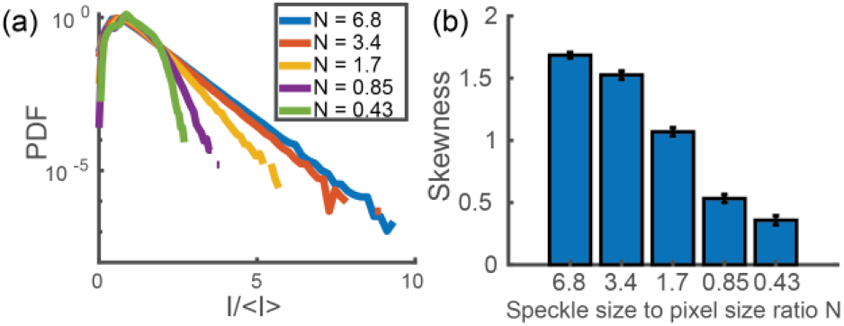
Sampling and integrity of the intensity distribution. The ratio of mean speckle width to the pixel width (N) plays an important role in determining whether speckles are properly measured. We synthetically downsampled pixels in a speckle pattern from the dynamic spatial data to reduce the speckle-to-pixel size ratio. Histograms of different speckle-to-pixel size ratios (a) and the skew plot (b) show that only if the speckles are sampled highly enough will the intensity distribution be adequately represented, with skewness reduced with inadequate sampling.

In conclusion, this study demonstrates the feasibility of measuring full speckle statistics through MMF bundles using a SCOS/T methodology. We confirmed that the granular speckle patterns arising from coherent interference can be detected through MMFs. Statistics of the speckle intensity distribution after traveling through MMFs still differentiate speckle from dark and system noise and relay flow information. In summary, multi-mode fiber bundles are a promising approach to developing both SCOS and SCOT systems for measuring CBF in humans and the results herein place these methods on a more secure statistical and analytical foundation.

## Data availability

Data underlying the results presented in this paper are not publicly available at this time but may be obtained from the authors upon reasonable request.

